# Perceptual Invariance of Words and Other Learned Sounds in Non-human Primates

**DOI:** 10.1101/805218

**Authors:** Jonathan Melchor, Isaac Morán, Tonatiuh Figueroa, Luis Lemus

## Abstract

The ability to invariably identify spoken words and other naturalistic sounds in different temporal modulations and timbres requires perceptual tolerance to numerous acoustic variations. However, the mechanisms by which auditory information is perceived to be invariant are poorly understood, and no study has explicitly tested the perceptual constancy skills of nonhuman primates. We investigated the ability of two trained rhesus monkeys to learn and then recognize multiple sounds that included multisyllabic words. Importantly, we tested their ability to group unexperienced sounds into corresponding categories. We found that the monkeys adequately categorized sounds whose formants were at close Euclidean distance to the learned sounds. Our results indicate that macaques can attend and memorize complex sounds such as words. This ability was not studied or reported before and can be used to study the neuronal mechanisms underlying auditory perception.

## Introduction

The ability to recognize the identity of a sound through variations in sensory input, such as a specific vocalization emitted by different talkers, exists in humans and likely in other animals(Elie & Theunissen, 2015; Peterson & Barney, 1952; Saunders & Wehr, 2019; Seyfarth, Cheney, & Marler, 1980; Town, Wood, & Bizley, 2018). Although this ability is vital for communication in primates, the perceptual basis of invariant recognition of sounds has been scarcely investigated. One possible reason for this is that non-human primates may only show limited acoustic learning(Fritz, Mishkin, & Saunders, 2005; Scott, Mishkin, & Yin, 2012; Wright, 1999), so their recognition capability may depend on genetically-programmed circuits(Brockelman & Schilling, 1984; Owren, Dieter, Seyfarth, & Cheney, 1992; Zador, 2019). On the other hand, it is known that macaques are capable of learning repertoires of visual categories(Rajalingham, Schmidt, & DiCarlo, 2015) and report the existence of objects with ambiguous or incomplete information(Diamond et al., 2016; Roy, Buschman, & Miller, 2014). However, this ability has never been tested for acoustic perception in non-human primates. In this paper, we sought to determine what acoustic parameters drive the invariant recognition of sounds (IRS) in trained non-human primates. We hypothesized that monkeys would invariably recognize sounds of salient patterns that resembled those the animals learned(Furuyama, Kobayasi, & Riquimaroux, 2017; Remez, Rubin, Pisoni, & Carrell, 1981). To further test this, we designed a novel paradigm in which the macaques had to report the recognition of target (T) sounds presented in sequences that included nontarget (N) sounds. We found that the monkeys invariantly recognized unexperienced sounds of frequency patterns near prominent patterns of learned sounds. Our results allowed us to elucidate the acoustic parameters(Furuyama, Kobayasi, & Riquimaroux, 2016; Ghazanfar et al., 2007; Shue, Keating, & Vicenik, 2009; Tchernichovski, Nottebohm, Ho, Pesaran, & Mitra, 2000) that lead to monkeys’ IRS. We also demonstrate that rhesus monkeys are capable to learn diverse sounds of complex spectrotemporal structures such as words. In addition, we demonstrate that the monkeys perceive unheard versions to be invariant of the related learned categories.

## Results

In order to study the invariant recognition of sounds, we trained two rhesus monkeys in an acoustic recognition task. During the task, the monkeys obtained a reward for releasing a lever after identifying a T presented after zero, one or two Ns (Fig. 1a-c; see Methods). After two years of training the monkeys learned to guide their behaviour attending acoustic information. Since then, the monkeys included numerous sounds into T or N categories by discovering, in few trials, which delivered reward and which did not. Then, the monkeys consolidated their memories by practicing few sounds during several days, and when their behaviour was consistent, we delivered new sounds. This phase of training took no more than two months, and then we decided to limit the number of sounds the monkeys would learn in order to privilege the number of repetitions per sound for each sound during the experiments. Overall, monkey V recognised seven Ts and twenty-one Ns, and monkey X eleven Ts and ten Ns. The macaques demonstrated excellent performance with an overall hit rate of 96.8 ± 0.11 (mean ± SEM, one-sample sign test, *p* < 0.01) (Supplementary Table 1). They also exhibited longer reaction times during false alarms (395.2 ± 128.4 ms) than during hits (281.8 ± 63.8 ms, Kruskal-Wallis test, *p* < 0.001) (Supplementary Fig. 1). Figs. 1d and 1e present examples of five Ts and five Ns frequently used during the experiments. The hit rate of monkey V was better when Ts in the first position, whereas monkey X was faster for Ts presented in the first and third positions (Kruskal-Wallis test, *p* < 0.01).

**Fig. 1.**
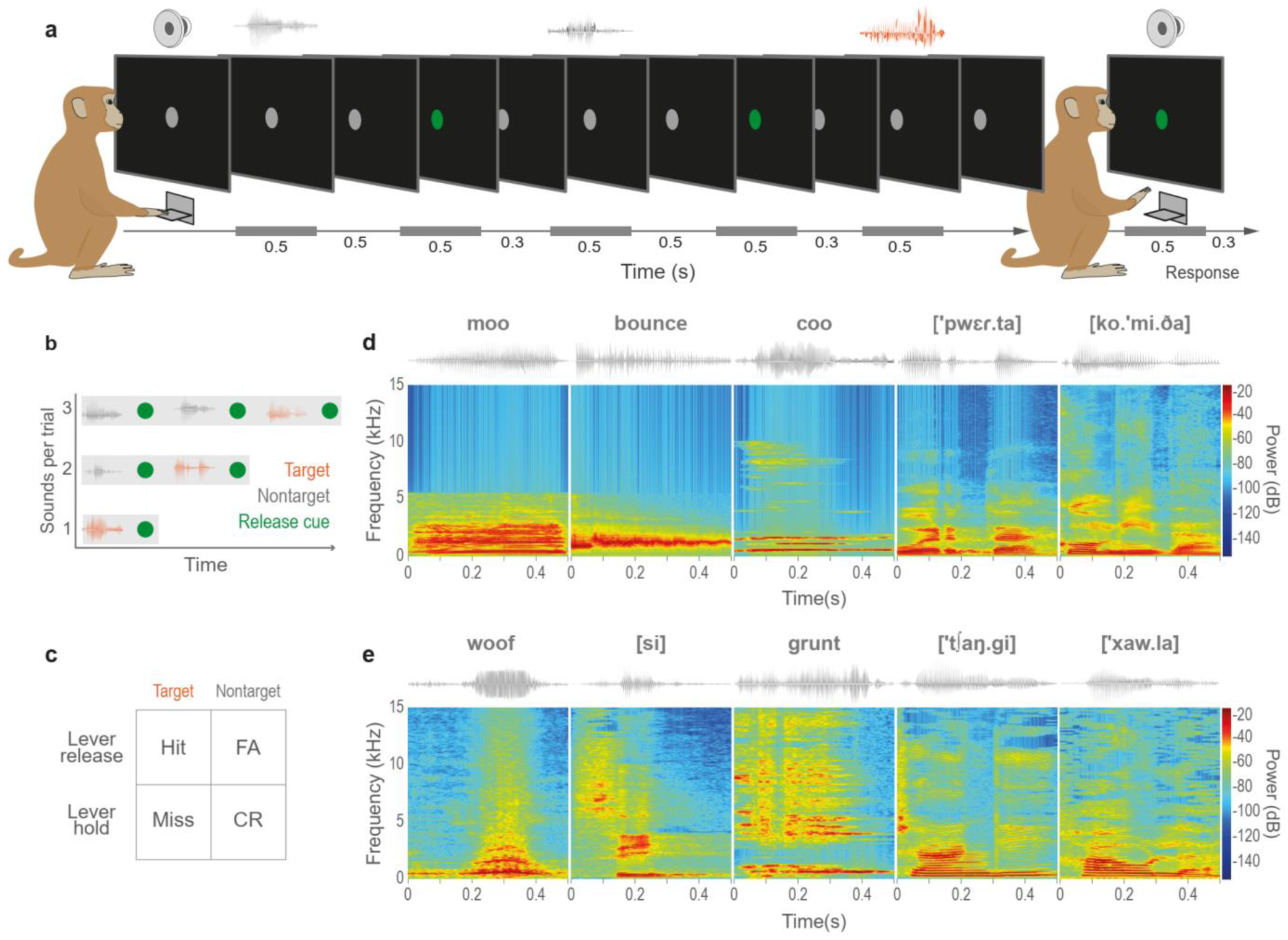
Auditory recognition task. **a** An example of the sequence of events of a trial. First, a visual cue appeared at the center of the screen to indicate that the monkey should press and hold the lever down. After a variable period of 0.5 to 1 s, a playback of 1 to 3 sounds commenced, each followed by a 0.5 s delay and a 0.5 s green cue (G). The monkey obtained a drop of liquid for releasing within 0.7 s of the beginning of the G that followed the T. Releases at other periods aborted the trial. Colour code: orange=T, grey=N, green=release cue. **b** Depictions of sequences of one, two or three sounds. Note that T always appeared last. **c** The behavioural outcomes after presentations of Ts and Ns. FA, false alarm, CR, correct rejection. **d** Sonograms and spectrograms of five Ts. IPA nomenclature describes Spanish words used in the experiments. **e** same as in (**d**) but for nontarget sounds.

### The monkeys recognized sounds based on their mean frequencies

To study the monkeys’ ability to differentiate Ts from Ns, we inquired the monkeys with sets of morphed sounds created from mixtures of a T and an N in different proportions (see Methods). Fig. 2a illustrates a morphing set in which the N /si/ (i.e. the Spanish word for ‘yes’) gradually morphed into a T coo monkey call. Fig. 2b shows psychometric functions (PFs) of the probability of recognising a morph as a T. Here, the differential limen (DL) indicates the minimum proportion of T required for recognition of a morph. There were no differences between monkey V’s and monkey X’s DLs: 11.3 ± 1.2 and 10.93 ± 1.4 (mean ± SEM), respectively (Kruskal-Wallis test, *p* = 0.93; Supplementary Table 2). In order to elucidate the acoustic variables responsible for recognitions, we calculated acoustic functions of morph parameters (e.g. AM, periodicity, entropy and pitch; see Methods) to contrast to the PFs. Thus, we derived Pearson acoustic functions (PAFs) from Pearson correlations of each morph and 100% T (Fig. 2c). Therefore, the PAFs express the similarities between the morphs’ acoustic modulations and the modulation of T. Nevertheless, as an alternative, we computed acoustic functions of the Euclidean distances (FEDs) between parameters in the morphs and in T (Fig. 2d). Finally, to determine whether recognition of morphs as T depended on Pearson or on proximities to acoustic parameters, we performed Spearman correlations of PAFs and FEDs with the PFs (Figs. 2e and 2f, respectively). The results indicated that FEDs of mean frequencies were strongly correlated with performance. Here, the average rho-values were 0.97 and 0.96 for monkeys V and X, respectively (*p* < 0.05, Supplementary Table 3), meaning that acoustic saliencies such as the formants drove the monkeys’ abilities to recognise sounds.

**Fig. 2.**
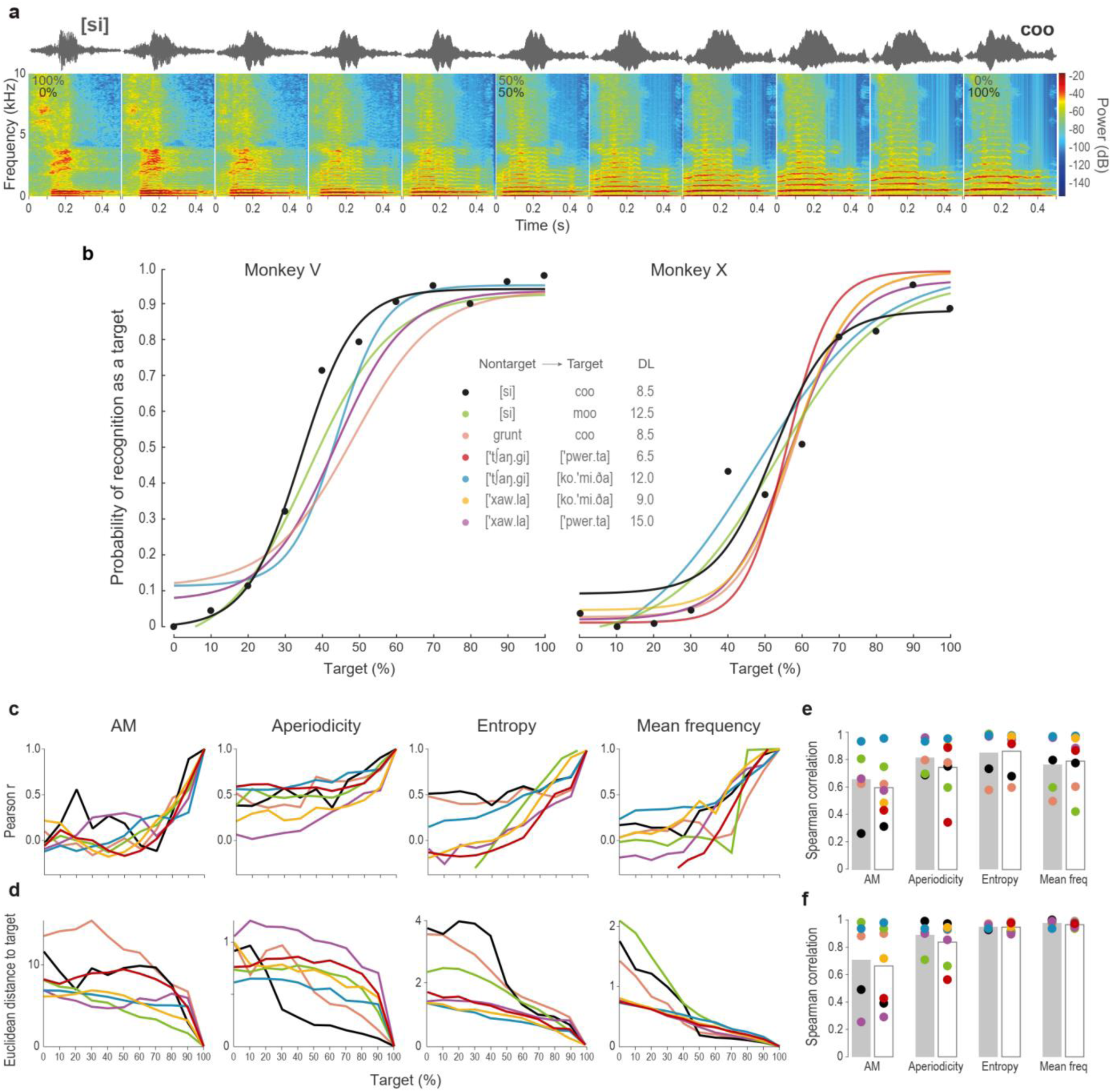
The mean frequency correlates to target recognitions. **a** An example of a morphing set in which a N [si] morphed to an T coo, from 0% T to 100% T in increments of 10%. Every morphing set comprised eleven morphs. **b** Monkey V’s and monkey X’s probabilities of recognising a morph as a T during the morphing set shown in (**a**). Continuous lines correspond to the sigmoidal fit to the average performance during the different morphing sets. **c** Subpanels present Pearson’s acoustic functions (PAFs) of various acoustic metrics (see Methods). Same colours as in (**b**). **d** Same as in c but for acoustic functions of Euclidean distances (FEDs). **e** Each dot is a Spearman correlation coefficient (rho) between the psychometric functions and PAFs, for different acoustic metrics. Same colours as in previous panels. Solid bars, monkey V. Unfilled bars, monkey X. **f** Same as in (**e**) but for FEDs.

### Invariant recognition arises from variants at Euclidean proximities to learned sounds

To test for IRS in macaques, we presented the monkeys with several versions of the learned sounds, e.g. one word uttered by different individuals. We experimented with sets of five versions of each T and N. Fig. 3a presents the T [‘pwεɾ.ta] spectrogram, i.e. the Spanish disyllabic word for door, and five variants (v1-v5). The boxplots in Fig. 3b correspond to the probabilities of recognising the versions as a T. The monkeys recognised 78.0% of the fifty versions above chance (one-sample sign test, *p* ≤ 0.05), with no performance differences between the two monkeys: 84.4 and 84.3% hit rate (Mann-Whitney test, *p* = 0.148). To determine whether the recognition of a version was due to the Euclidean proximity between any of its acoustic parameters to a learned sound, we calculated various FEDs from various acoustic parameters. Fig. 3c shows that, using the parameter ‘Mean Frequency’, the Euclidean distances of [‘pwεɾ.ta] to four of its versions were smaller than the distances of those versions to other learned sounds. The only exception was a version closer to the coo sound. However, the normalised distances showed that the version of [‘pwεɾ.ta] closer to the coo produced the lowest performance (Fig. 3d). Similarly, Fig. 3e shows that the mean frequency of variants of other learned sounds were also closer to the expected category (Spearman correlation, R = 0.92, *p* < 0.01). Moreover, the FEDs of the sounds’ mean frequencies explained performance better than PAFs and other acoustic parameters (Fig. 3f).

**Fig. 3.**
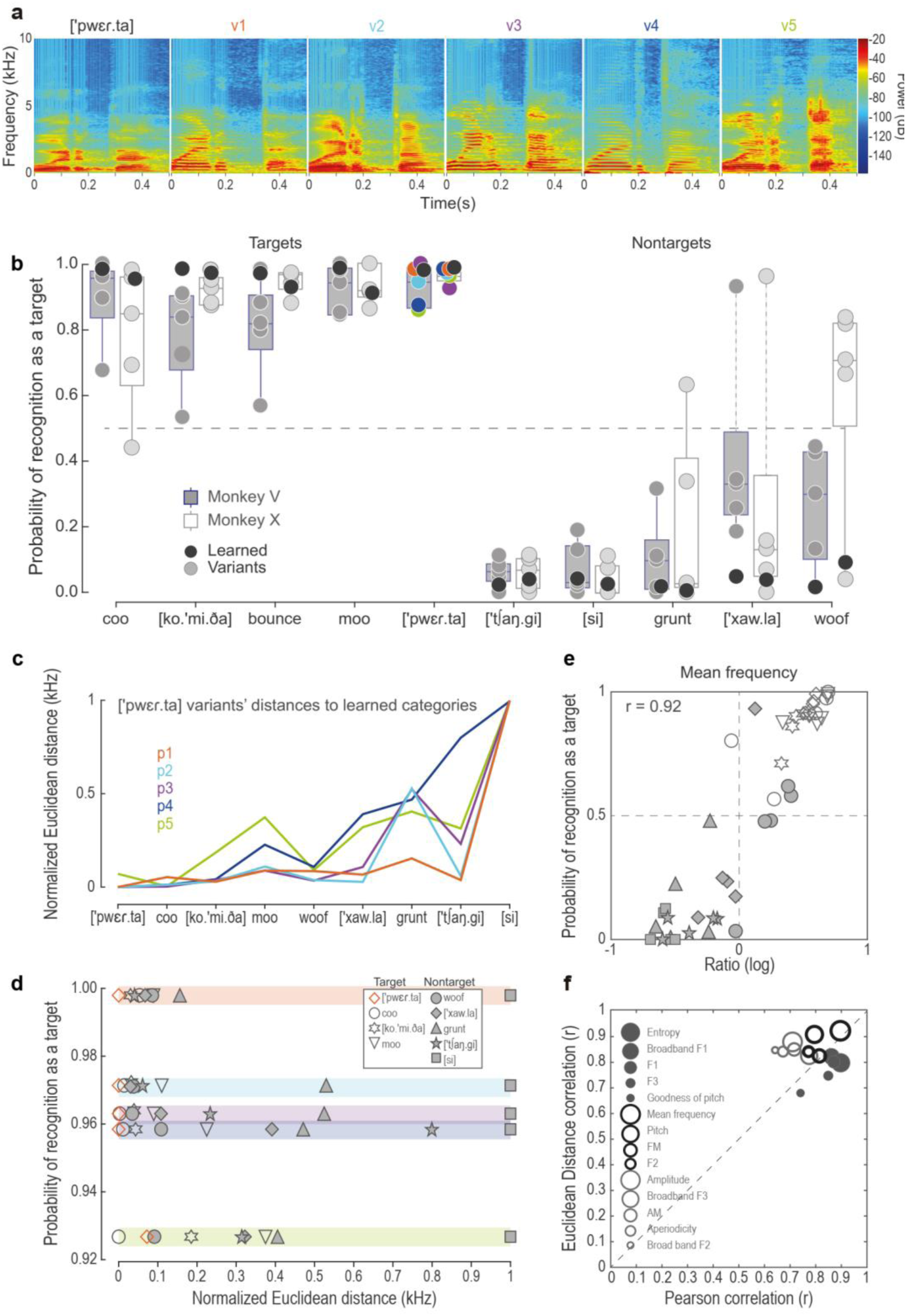
Mean frequency proximities between learned sounds and their variants produce perceptual invariance. **a** Spectrograms of T [‘pwer.ta], i.e. the Spanish word for door and five variants. Each variant corresponds to a different speaker (v1-v5). **b** Boxplots of the probability of recognising a variant as a T. Colours at [‘pwer.ta] categories correspond to variants at (**a**). **c** Normalised Euclidean distances of variants of [‘pwer.ta] to four Ts and five Ns. Colours are the same as in (**a**) and (**b**). Symbols are labelled at the abscissas. **d** Mean monkey performance as a function of Euclidean distances of variants of [‘pwer.ta] to all Ts and Ns. Same colour code as in (**a-c**). **e** Logarithmic ratio of the recognition of variants at (**b**) and the mean-frequency distance to each T and N, plotted as a function of the probability of recognising a T. Symbols as in (**d**). Upper left, the Pearson correlation coefficient (r) between the logarithmic ratio and behaviour. **f** Similar to (**e**) but for Pearson’s r, and for all of the tested acoustic metrics.

### The formants of the sounds contribute to IRS

Since the mean frequency is derived from the mean power of the frequencies in a sound, we explored the contribution to IRS of the frequencies with highest power modulations, e.g. the acoustic formants. To do this, we presented the monkeys with sounds of some formants of the learned sounds and their versions. Fig. 4a shows spectrograms of the T [ko.’mi.ða], i.e. the Spanish trisyllabic word for food, and its F1, F2, and F1&F2 formants. Similarly, Fig. 4b shows spectrograms of a version of [ko.’mi.ða], and its formants. The hypothesis was that formants of the learned sounds would suffice to drive the monkeys’ recognitions. Moreover, that formants of the versions modulated in the range of the learned sounds would also work for acoustic recognition (Fig. 4c). The monkeys performed for no more than forty presentations of each sound in order to prevent the learning of formants as T or N. Fig. 4d presents the mean performance of the monkeys during the recognition of sounds in Fig. 4a-b.

**Fig. 4.**
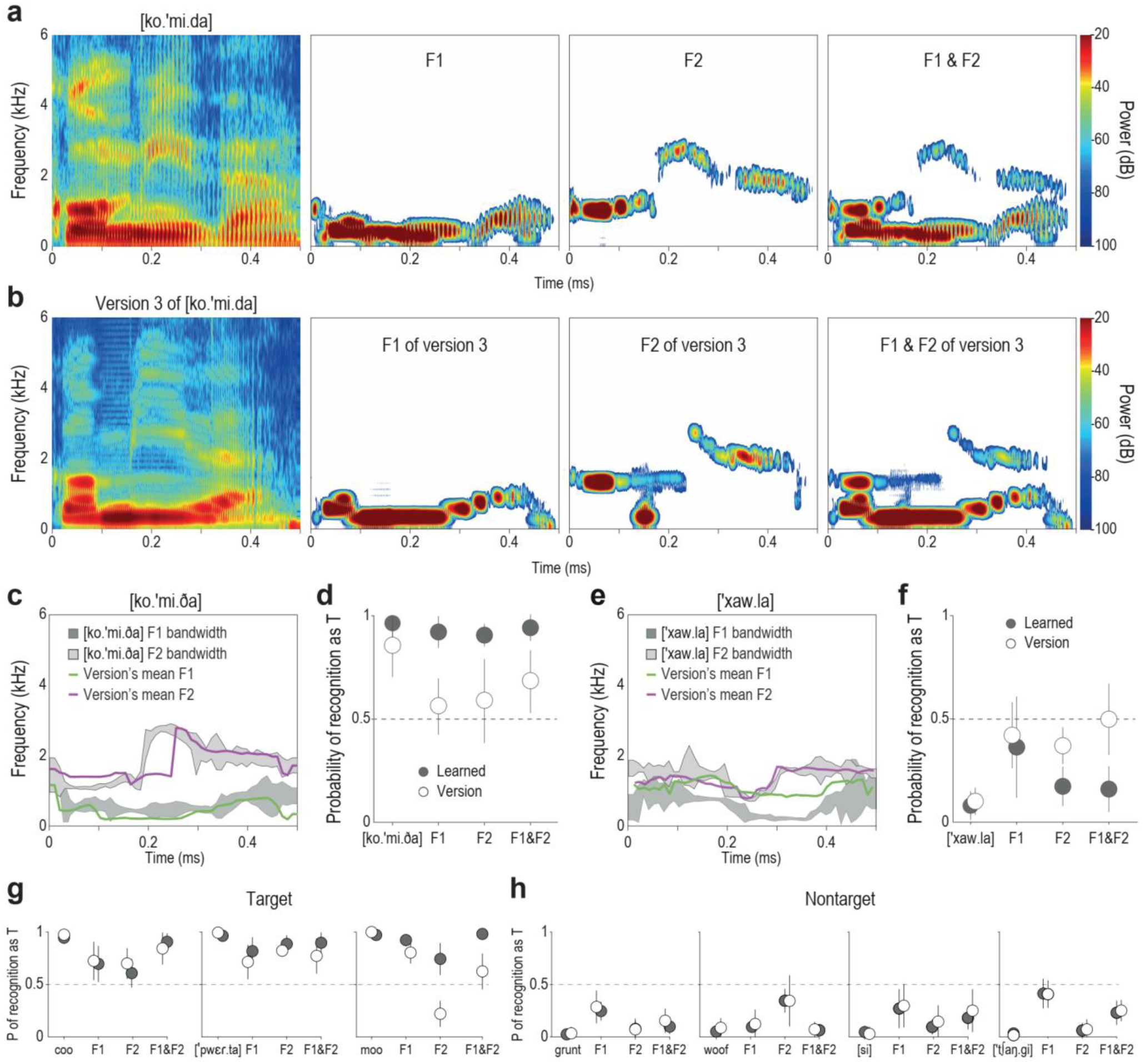
The first and second formants are key for perceptual invariance. **a** Spectrograms of T [ko.’mi.da], and first, second, and first & second formants. **b** One version of [ko.’mi.da], and its corresponding formants. **c** Comparison of F1 and F2 bandwidth formants of [ko.’mi.da] and the mean of the version’s F1 and F2 formants. **d** Monkeys’ mean probability of recognising sounds in (a) as T. **e-f** Same as in (**c**) and (**d**) but for N [‘xaw.la]. **g-h**, Probability of recognition for F1, F2, F1&F2, learned and variants of Ts and Ns, respectively.

The monkeys significantly identified [ko.’mi.ða], its formants, the versions, and the versions’ F1&F2 formants (one-sample sign test, *p* < 0.01). However, the versions’ F1 or F2 alone were not sufficient for recognition. Fig. 4e is the same as Fig. 4c but for the category [‘xaw.la]. Fig. 4f shows false alarms of [‘xaw.la], the versions and formants. Here, F2 of the learned and version sounds, and F1&F2 of the learned sound did not produce a significant number of false alarms. Finally, Figs. 4g-h present the results for other Ts and Ns, and their versions. The monkeys recognised the learned T with a probability of 0.93 ± 0.03, and versions of Ts with a P of 0.73 ± 0.09. Meanwhile, the false alarms of learned N had a P of 0.14 ± 0.06, and for Ns versions P was 0.24 ± 0.16. Overall, 94% of F1&F2 of learned and version sounds were recognised significantly (one-sample sign test, *p* < 0.01) (Supplementary Table 4). These results suggest that the invariant recognition of sounds in macaques is created from acoustic saliencies modulated in the range of saliencies of learned sounds.

## Discussion

We presented evidence of the invariant recognition of sounds in monkeys. This evidence is mainly supported by the ability of the monkeys to recognise variants to which they had no previous exposure. The learned sounds included words and naturalistic sounds in a broad range of frequencies and temporal modulations. Remarkably, the recognition of the variants was based on their Euclidean proximity to the saliences of the learned sounds. To our knowledge, this is the first demonstration of the ability of monkeys to store in long-term memories information about the sound of words and other naturalistic tokens.

### Macaques learn numerous naturalistic sounds

The training of monkeys was indeed more tenuous and prolonged than in visual or tactile paradigms(Lemus, Hernández, & Romo, 2009; Rajalingham et al., 2015) but achievable, they recognised sounds that included multisyllabic words above a hit rate of 90%. This single result suggests that acoustic circuits cannot be entirely based on genetic programmes(Brockelman & Schilling, 1984; Owren et al., 1992; Zador, 2019), similar to recently reported in songbirds(Moore & Woolley, 2019). Moreover, we verified that the learned sounds remained in long-term memories because the monkeys were able to solve the task effective after periods of up to five weeks of rest.

A realistic possibility was that the monkeys only learned the first or the last chunks of the sounds. Nevertheless, since the macaques had to wait for 0.5 s after each sound to respond they probably accumulated all available evidence, similar to previous reports showing that they needed all disposable information for discriminate acoustic flutter-frequencies(Lemus et al., 2009), for example. A weakness of our study was the lack of semantic relationships to each of the sounds. Perhaps with the only exception of the conspecific vocalizations, other sounds have no particular meaning for the monkeys other than being T or N. If this was the case, it is interesting to note that the monkey vocalizations acquired and alternative meaning to the monkeys; i.e., T or N, which also mean reward and holding down the lever, respectively. Nevertheless, in our study, the repertoire of frequencies within the Ts and Ns were likely to form diverse neural representations throughout the superior temporal gyrus. Similar associations to behaviour may occur in other communicating animals(Elie & Theunissen, 2015; Saunders & Wehr, 2019; Town et al., 2018).

### Acoustic recognition arises from a rule of proximity

To understand the IRS, it is fundamental to discern the range of acoustic variability where a perceptual category remains. Our first hypothesis was that the IRS emerged from the similarity of acoustic modulations between a learned sound and its versions. Thus, we first searched for Pearson’s correlations between the continuous functions of the learned sounds and the mixtures of T and N categories. The relationships would suggest the existence of spectrotemporal fingerprints emulated by the morphs. However, we found that subtle differences ruled out the hypothesis. Alternatively, we found that the perceptual constancy of acoustic categories occurred for versions with mean frequencies at short Euclidean distances of the learned sounds. This finding coincides with recent reports on vowel identification(Town et al., 2018), and is consistent with the notion of formants being crucial for carrying acoustic identities(Fitch & Fritz, 2006; Furuyama et al., 2016, 2017; Ghazanfar et al., 2007; Remez et al., 1981). One explanation is that the salient formants emerge with semantic information from the persistent fine structure of sounds, such as timbre, which may be responsible for streaming —as in a cocktail party paradigm. In such a scenario, perhaps neuronal responses that adapt to timbre code only for the formants. One possible consequence would be that speakers learn to modulate formants in order to communicate, and not the pitch, nor the timbre, which are more useful for sound localisation, or recognition of conspecifics(Takahashi, Fenley, & Ghazanfar, 2016). This possibility, however, needs to be corroborated in future experiments, where experimental models as the one we present here, may become crucial.

### Hierarchical processing of sounds

Recordings of neurons in passive untrained macaque have demonstrated that belt area neurons around the core of the auditory cortex (A1) are responsive to band-passed noises(Rauschecker & Tian, 2004), FM-sweeps(Biao Tian & Rauschecker, 2004), and conspecific vocalisations(Ortiz-Rios et al., 2017; Rauschecker & Tian, 2000; B. Tian, Reser, Durham, Kustov, & Rauschecker, 2001). The belt receives the information contained in vocalisations from simultaneous projections of the neurons which demonstrate sharp frequency tuning in A1. However, since these cells also respond to reversed monkey calls(Recanzone, 2008), they do not code for specific sequences of frequencies that provide identity to the acoustic categories. This would suggest and support the finding of PFC neurons encoding for vocalisations organised in specific frequency sequences(Cohen, Hauser, & Russ, 2006; Romanski, Averbeck, & Diltz, 2005; Russ, Ackelson, Baker, & Cohen, 2008). Nevertheless, those cells were observed in non-behaving macaques, so their contribution to acoustic perception remains unclear. In order to understand what parameters correlate with auditory perception, experiments using monkeys trained to discriminate the syllables /bad/ and /dad/ found categorical responses to linear mixtures of the syllables at the belt(Tsunada, Lee, & Cohen, 2011). This finding means that belt neurons responded to perceptual categories and not to particular spectrotemporal modulations. Recent fMRI studies in humans and macaques showed that anterior areas of the superior temporal gyrus respond more to conspecific vocalisations compared to other sounds(Leaver & Rauschecker, 2010; Perrodin, Kayser, Abel, Logothetis, & Petkov, 2015; Perrodin, Kayser, Logothetis, & Petkov, 2011; Petkov et al., 2008; Robert J. Zatorre; Pascal Belin, 2001; Shue et al., 2009), suggesting a distributed cortical representation of sounds relevant to behaviour. An important question is whether those representations serve as templates for the recognition of similar sounds(Belin, Bodin, & Aglieri, 2018). Studies of the inferotemporal and prefrontal cortices of monkeys showed neurons whose categorical responses achieved the grouping of wide variations of images(Bao & Tsao, 2018; DiCarlo, Zoccolan, & Rust, 2012; Seger & Miller, 2010), consistently with perceptual reports(Cromer, Roy, & Miller, 2010; Wutz, Loonis, Roy, Donoghue, & Miller, 2018). Similarly, experiments in the prefrontal cortex and secondary auditory areas suggest the neuronal coding of acoustic categories(Cohen et al., 2006; Leaver & Rauschecker, 2010; Perrodin et al., 2015, 2011; Petkov et al., 2008; Romanski et al., 2005; Russ et al., 2008; Tsunada et al., 2011). Experiments conducted with behaving ferrets showed that A1 neurons can respond to variations of vowels(Town et al., 2018). However, the neurons were sensitive to input timing, suggesting that the recognition of longer and more complex sounds requires further cortical integration.

Based on our results, it’s probably that recognition circuits hierarchically integrate patterns of acoustic prominences, including combinations, as in words. Furthermore, recurrent sounds create neuronal templates, sometimes evoked by similar saliencies of variants. Further experiments may explore semantics using our auditory paradigm. For example, the coding of the meaning of conspecific vocalizations in different brain areas(Chandrasekaran, Lemus, & Ghazanfar, 2013; Ortiz-Rios et al., 2015; Petkov et al., 2008; Rauschecker & Tian, 2000; Rauschecker, Tian, & Hauser, 1995; Recanzone, 2008; Robert J. Zatorre; Pascal Belin, 2001; B. Tian et al., 2001). In conclusion, the behavioural paradigm we present could serve to advance the study of acoustic recognition at the neuronal level, because, in contrast to humans(Coupé, Oh, Dediu, & Pellegrino, 2019), trained monkeys present only a few dozen acoustic representations, meaning fewer lexical overlaps, which could benefits the study of discrete acoustic percepts.

## Methods

### Ethics statement

All procedures were performed in compliance with the Mexican Official Standard for the Care and Use of Laboratory Animals (NOM-062-ZOO-1999) and approved by the Internal Committee for the Use and Care of Laboratory Animals of the Institute of Cell Physiology, UNAM (CICUAL; LLS80-16).

### Animals and experimental setup

Two adult rhesus macaques (*Macaca mulatta*; one male, 13 kg, ten yrs. old, and one female, 6 kg, ten yrs. old) participated in this study. Typically, each monkey performed ∼1000 trials during sessions of three hours (one session per day, six sessions per week). The monkeys received a daily minimum water intake of 20 ml/kg, completed in cage as needed. The monkeys’ training lasted approximately two years and concluded after each one recognised more than 20 sounds above an ∼85% hit rate. Training and experimental sessions took place in a soundproof booth. The macaque was seated in a primate chair, 60 cm away from a 21” LCD colour monitor (1920 × 1080 resolution, 60 Hz refreshing rate). A Yamaha MSP5 speaker (50 Hz – 40 kHz frequency range) was placed fifteen cm above and behind the monitor to deliver acoustic stimuli at ∼65 dB SPL (measured at the monkeys’ ear level). Additionally, a Logitech® Z120 speaker was situated directly below the Yamaha speaker in order to render background white noise at ∼55 dB SPL. Finally, a metal spring-lever situated at the monkeys’ waist level captured the responses.

### Behavioural Task

The acoustic recognition task (ART) consisted of identifying T and N sounds. Fig. 1a presents the elements of the paradigm as follows: First, a grey circle with an aperture of 3° appeared at the centre of the screen, and the monkey pressed and held down the lever. Immediately thereafter, a playback of from 1 to 3 sounds began, and a T was always the last sound (Fig. 1b). After each sound, the monkey kept the lever down for another 0.5 s until the visual cue turned green (G). If the audio was a T, the monkey had 0.8 s to release the lever and receive a drop of liquid. However, releases at other periods constituted a false alarm (FA) that led to the abortion of the trial (Fig. 1c). The task’s programming was in LabVIEW 2014 (SP1 64-bits, National Instruments®).

### Stimuli

The sounds were recordings from our laboratory or downloads from free internet libraries. They consisted of natural and artificial environmental sounds, e.g. monkey calls, other animal vocalisations and words. All sounds were sampled at 44.1 kHz (cutoff frequencies: 100 Hz to 20 kHz), amplitudes were normalised at −10 dB SPL (RMS), and compressed or elongated to 0.5 s. Fig. 1d presents examples of five T and five N used frequently during the experiments. The morphing sets comprised 11 mixtures of T and an N in proportions ranging from 0% T (i.e. 100% N) to 100% T in 10% increments of T(Chakladar, Logothetis, & Petkov, 2008; Kawahara, Masuda-Katsuse, & De Cheveigné, 1999). Each morphed sound was repeated randomly ten times but always presented first in a trial. Trials of two or three sounds were completed with T and N. To test for IRS, versions of learned sounds were presented forty times randomly, but only after the monkeys’ training concluded. Finally, we examined the recognition of acoustic salience using F1, F2, and F1&F2 formants of learned sounds and versions. All sounds were processed using Adobe Audition® version 6.0. The morphed sounds were created using the signal processing software STRAIGHT(Kawahara et al., 1999) (Speech Transformation and Representation based on Adaptive Interpolation of Weighted spectrograms: http://www.wakayama-u.ac.jp/~kawahara/STRAIGHTadv/index_e).

### Analysis

PFs were TanH regressions of the probability of recognising a morph as a T(Duarte, Figueroa, & Lemus, 2018; Duarte & Lemus, 2017). PAFs and FEDs were functions of Pearson correlations between continuous parameters measured at each morph and the same parameter in 100% T, and the Euclidean distances from each M to 100% T, respectively(Town et al., 2018). Spearman correlations between FAP and FED with PF computed the contribution of acoustic parameters to recognition. Differential limen (DL) was half the difference between the abscissa projected to the PF at 75%, and 25% performance. Reaction times were times of lever releases after the start of G. Logarithmic ratio = log(performance) – log(distance). Behavioural analyses were performed using SigmaPlot® version 12.0 software for Windows (Systat Software, Inc., San Jose, CA, USA), and customised algorithms in MATLAB® 8.5.0.1, R2015a (The Mathworks, Inc). Acoustic metrics were computed using Pratt (Boersma, P., & Van Heuven, 2001) (version 6.0.37, http://www.fon.hum.uva.nl/praat/), VoiceSauce (Shue et al., 2009)(version 1.36, http://www.seas.ucla.edu/spapl/voicesauce/) and Sound Analysis Pro(Tchernichovski et al., 2000) (http://soundanalysispro.com/).

## Notes

#### Summary of Updates

The description of the training of the monkeys was improved, and supplementary information simplified. Additionally, the introduction now has only the most relevant information.

