## Supplementary material for "Perceptual Invariance of Words and Other Learned Sounds in Non-human Primates": Supplemmentary Figure 1

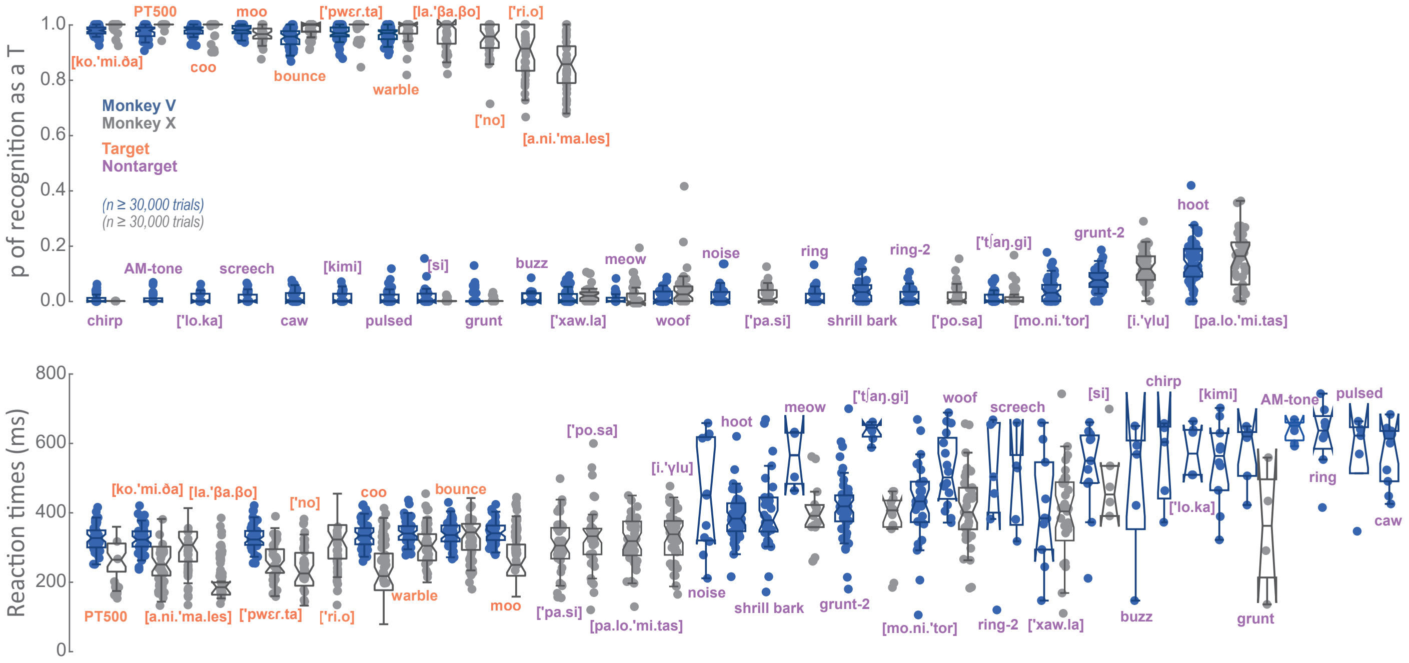


**Supplementary Figure 1.** Behavioural performance and reaction times (RT). Probability of recognition (top) and RT (bottom) for all learned target (orange) and nontarget (purple) sounds by the monkeys V (blue) and X (gray). All  the sounds were 0.5 s in duration.
