## Supplementary material for "Perceptual Invariance of Words and Other Learned Sounds in Non-human Primates": Table 1

**Supplementary Table 1. Probability of recognition and description of the sound set.**

| | Name | Description | P of recognition as T | |
|---|---|---|---|---|
| | | | Monkey V | Monkey X |
| Target | coo | Rhesus monkey vocalization | 97.76 ± 02.04 | 98.90 ± 02.82 |
| | warble | Rhesus monkey vocalization | 96.33 ± 02.55 | 97.81 ± 03.99 |
| | [ko.'mi.ða] | Spanish word for food | 97.59 ± 01.70 | 99.37 ± 01.76 |
| | ['pwɛɾ.ta] | Spanish word for door | 96.58 ± 02.85 | 98.80 ± 03.06 |
| | [a.ni.'ma.les] | Spanish word for animals | - | 84.97 ± 09.31 |
| | ['ři.o] | Spanish word for river | - | 89.91 ± 08.84 |
| | [no] | Spanish word for not | - | 94.86 ± 05.97 |
| | [la.'βa.βo] | Spanish word for sink | - | 96.40 ± 05.16 |
| | moo | Vocal sound of a cow | 97.93 ± 01.88 | 96.34 ± 02.75 |
| | bounce | Bouncing tone | 95.01 ± 03.23 | 98.31 ± 02.54 |
| | PT500 | Pure tone of 500 Hz | 97.09 ± 02.23 | 99.24 ± 01.85 |
| Nontarget | grunt | Rhesus monkey vocalization | 01.26 ± 02.29 | 00.19 ± 00.67 |
| | grunt-2 | Rhesus monkey vocalization | 07.78 ± 04.20 | - |
| | shrill bark | Rhesus monkey vocalization | 03.95 ± 04.06 | - |
| | pulsed | Rhesus monkey vocalization | 01.31 ± 02.55 | - |
| | ['lo.ka] | Spanish word for crazy | 00.97 ± 01.67 | - |
| | [kimi] | Spanish pseudoword | 01.07 ± 01.88 | - |
| | ['tʃaŋ.gi] | Spanish pseudoword | 01.22 ± 02.63 | 01.58 ± 03.42 |
| | [si] | Spanish word for yes | 01.37 ± 02.81 | 00.18 ± 00.54 |
| | ['xaw.la] | Spanish word for cage | 01.42 ± 02.23 | 02.50 ± 03.13 |
| | [mo.ni.'tor] | Spanish word for computer monitor | 03.91 ± 04.31 | - |
| | [pa.xa.'ri.tos] | Spanish word for birdies | - | 00.00 ± 00.00 |
| | ['po.sa] | Spanish word for pose | - | 02.13 ± 03.51 |
| | ['pa.si] | Spanish pseudoword | - | 02.33 ± 03.00 |
| | [i.'ɣlu] | Spanish word for igloo | - | 12.27 ± 06.06 |
| | [pa.lo.'mi.tas] | Spanish word for popcorn | - | 15.31 ± 09.66 |
| | meow | Cat meowing | 00.90 ± 01.76 | 02.39 ± 04.48 |
| | chirp | Bird vocalization | 00.92 ± 01.69 | - |
| | screech | Parrot screech | 01.01 ± 01.79 | - |
| | caw | Crow squawk | 01.57 ± 02.60 | - |
| | woof | Dog bark | 01.83 ± 02.34 | 04.51 ± 07.53 |
| | hoot | Owl hooting | 14.29 ± 07.95 | - |
| | AM-tone | Amplitude-Modulated tone (1 kHz) | 00.85 ± 01.66 | - |
| | buzz | Mosquito whine | 01.10 ± 01.82 | - |
| | ring | Cell phone ring tone | 01.41 ± 02.66 | - |
| | ring-2 | Ring bell | 01.68 ± 02.53 | - |
| | noise | Passband noise (1-4 kHz) | 01.87 ± 03.17 | - |

Data are mean ± SD; ($n \geq 41$ sessions).
The IPA nomenclature was obtained with the Spanish automatic phonetic transcriptionist, created for Xavier López Morrás (http://www.aucel.com/pln/transbase.html).
