## Supplemental Data 1 for "Perceptual Invariance of Words and Other Learned Sounds in Non-human Primates"

**Supplementary Table 2. Differential limen (DL) and sigmoidal fit parameters for the morphing sets.**

| | Parameters of *tanh* function | Target and nontarget sounds for each morphing | | | | | | |
|---|---|---|---|---|---|---|---|---|
| | | [si] to coo | [si] to moo | grunt to coo | ['tʃaŋ.gi] to ['pwɛr.ta] | ['tʃaŋ.gi] to [ko.'mi.ða] | ['xaw.la] to [ko.'mi.ða] | ['xaw.la] to ['pwɛr.ta] |
| **Monkey V** | c | 0.47 | 0.44 | 0.53 | - | 0.51 | - | 0.53 |
| | α | 0.47 | 0.49 | 0.42 | - | 0.43 | - | 0.42 |
| | θ | 33.7 | 35.9 | 43.5 | - | 43.1 | - | 47.7 |
| | β | 0.07 | 0.05 | 0.08 | - | 0.05 | - | 0.05 |
| | Q | 0.65 | 0.65 | 0.46 | - | 0.62 | - | 0.46 |
| | DL | 8.50 | 12.5 | 8.50 | - | 12.0 | - | 15.0 |
| **Monkey X** | c | 0.47 | 0.49 | 0.45 | 0.50 | 0.49 | 0.52 | 0.51 |
| | α | 0.50 | 0.40 | 0.55 | 0.49 | 0.47 | 0.47 | 0.48 |
| | θ | 52.6 | 51.7 | 47.0 | 55.3 | 56.2 | 57.3 | 57.3 |
| | β | 0.04 | 0.07 | 0.03 | 0.09 | 0.06 | 0.07 | 0.07 |
| | Q | 0.49 | 0.44 | 0.61 | 0.80 | 0.75 | 0.65 | 0.70 |
| | DL | 15.5 | 11.0 | 16.5 | 6.50 | 9.50 | 9.00 | 8.50 |
