## Supplementary material for "Perceptual Invariance of Words and Other Learned Sounds in Non-human Primates": Table 2

**Supplementary Table 3.** **Spearmans' coefficients (Rho) between psychometric functions (PF) and Pearson Acoustic Function (PAF), and PF and Function of Euclidean distance (FED) for metrics.**

| | Rho from PF and PAF | | | Rho from PF and FED | | |
|---|---|---|---|---|---|---|
| **Acoustic metric** | **Monkey V** (n = 5) | **Monkey X** (n = 7) | **Proportion of Rho with p < 0.05** | **Monkey V** (n = 5) | **Monkey X** (n = 7) | **Proportion of Rho with p < 0.05** |
| AM | 0.66 ± 0.25 | 0.58 ± 0.21 | 58.30 | 0.70 ± 0.32 | 0.66 ± 0.28 | 58.30 |
| Aperiodicity | 0.81 ± 0.12 | 0.74 ± 0.21 | 83.30 | 0.88 ± 0.10 | 0.83 ± 0.16 | 91.70 |
| Entropy | 0.85 ± 0.18 | 0.86 ± 0.15 | 83.30 | 0.94 ± 0.01 | 0.95 ± 0.03 | 100.0 |
| Mean Frequency | 0.76 ± 0.21 | 0.78 ± 0.20 | 66.70 | 0.97 ± 0.02 | 0.96 ± 0.02 | 100.0 |
| Pitch | 0.93 ± 0.04 | 0.91 ± 0.06 | 100.0 | 0.94 ± 0.06 | 0.94 ± 0.04 | 100.0 |
| Amplitude | 0.92 ± 0.06 | 0.92 ± 0.03 | 100.0 | 0.95 ± 0.03 | 0.93 ± 0.02 | 100.0 |
| FM | 0.86 ± 0.08 | 0.86 ± 0.06 | 100.0 | 0.92 ± 0.05 | 0.90 ± 0.03 | 100.0 |
| Goodness of pitch | 0.78 ± 0.11 | 0.81 ± 0.13 | 91.70 | 0.84 ± 0.11 | 0.82 ± 0.11 | 100.0 |
| First formant | 0.86 ± 0.11 | 0.84 ± 0.16 | 91.70 | 0.94 ± 0.03 | 0.93 ± 0.03 | 100.0 |
| Second formant | 0.61 ± 0.35 | 0.56 ± 0.29 | 66.70 | 0.89 ± 0.03 | 0.77 ± 0.30 | 91.70 |
| Third formant | 0.76 ± 0.21 | 0.70 ± 0.19 | 75.00 | 0.78 ± 0.23 | 0.81 ± 0.14 | 83.30 |

Data are mean ± SD.
AM, Amplitude modulation; FM, Frequency modulation.
