## Supplementary material for "Perceptual Invariance of Words and Other Learned Sounds in Non-human Primates": Table 3

**Supplementary Table 4. Probability of recognition of learned and version sounds, and their formants.**

| Target (T) | Sound | | Formant frequency | | |
|---|---|---|---|---|---|
| | | | First (F1) | Second (F2) | F1 & F2 |
| | coo$_{learned}$ | 0.94 ± 0.05 | 0.69 ± 0.17 | 0.60 ± 0.13 | 0.90 ± 0.08 |
| | coo$_{version}$ | 0.97 ± 0.02 | 0.72 ± 0.18 | 0.70 ± 0.14 | 0.84 ± 0.15 |
| | moo$_{learned}$ | 0.97 ± 0.04 | 0.92 ± 0.05 | 0.74 ± 0.15 | 0.98 ± 0.03 |
| | moo$_{version}$ | 1.00 ± 0.00 | 0.80 ± 0.10 | 0.22 ± 0.12 (n.s.) | 0.62 ± 0.17 |
| | ['pwɛr.ta]$_{learned}$ | 0.96 ± 0.05 | 0.81 ± 0.13 | 0.88 ± 0.08 | 0.89 ± 0.09 |
| | ['pwɛr.ta]$_{version}$ | 0.99 ± 0.01 | 0.71 ± 0.16 | 0.82 ± 0.05 | 0.77 ± 0.16 |
| | [ko.'mi.ða]$_{learned}$ | 0.96 ± 0.05 | 0.92 ± 0.07 | 0.90 ± 0.05 | 0.94 ± 0.06 |
| | [ko.'mi.ða]$_{version}$ | 0.85 ± 0.12 | 0.56 ± 0.13 (n.s.) | 0.59 ± 0.20 (n.s.) | 0.68 ± 0.15 |
| | mean$_{learned}$ | 0.96 ± 0.01 | 0.84 ± 0.10 | 0.78 ± 0.13 | 0.93 ± 0.03 |
| | mean$_{version}$ | 0.95 ± 0.06 | 0.70 ± 0.10 | 0.58 ± 0.26 | 0.73 ± 0.09 |
| | Total | 0.95 ± 0.04 | 0.77 ± 0.12 | 0.68 ± 0.22 | 0.83 ± 0.12 |
| **Nontarget (N)** | | | | | |
| | grunt$_{learned}$ | 0.02 ± 0.04 | 0.24 ± 0.08 | 0.07 ± 0.09 | 0.09 ± 0.06 |
| | grunt$_{version}$ | 0.03 ± 0.05 | 0.28 ± 0.15 | 0.07 ± 0.08 | 0.15 ± 0.11 |
| | woof$_{learned}$ | 0.05 ± 0.05 | 0.09 ± 0.04 | 0.34 ± 0.11 | 0.06 ± 0.04 |
| | woof$_{version}$ | 0.08 ± 0.09 | 0.12 ± 0.13 | 0.34 ± 0.24 (n.s.) | 0.07 ± 0.07 |
| | [si]$_{learned}$ | 0.04 ± 0.06 | 0.27 ± 0.18 | 0.09 ± 0.07 | 0.18 ± 0.11 |
| | [si]$_{version}$ | 0.02 ± 0.06 | 0.29 ± 0.20 (n.s.) | 0.14 ± 0.15 | 0.25 ± 0.20 |
| | ['tʃaŋ.gi]$_{learned}$ | 0.03 ± 0.04 | 0.41 ± 0.14 (n.s.) | 0.06 ± 0.07 | 0.23 ± 0.11 |
| | ['tʃaŋ.gi]$_{version}$ | 0.01 ± 0.03 | 0.40 ± 0.13 (n.s.) | 0.07 ± 0.09 | 0.25 ± 0.10 |
| | ['xaw.la]$_{learned}$ | 0.07 ± 0.06 | 0.36 ± 0.24 (n.s.) | 0.17 ± 0.09 | 0.16 ± 0.10 |
| | ['xaw.la]$_{version}$ | 0.10 ± 0.06 | 0.42 ± 0.15 (n.s.) | 0.37 ± 0.08 | 0.49 ± 0.17 (n.s.) |
| | mean$_{learned}$ | 0.04 ± 0.02 | 0.27 ± 0.12 | 0.15 ± 0.11 | 0.14 ± 0.06 |
| | mean$_{version}$ | 0.05 ± 0.03 | 0.30 ± 0.12 | 0.20 ± 0.14 | 0.24 ± 0.16 |
| | Total | 0.05 ± 0.02 | 0.29 ± 0.11 | 0.17 ± 0.12 | 0.19 ± 0.12 |

**T & N** Proportion of sounds recognized significantly above chance level.

| | | | F1 | F2 | F1 & F2 |
|---|---|---|---|---|---|
| | Learned | 100 % | 77 % | 100 % | 100 % |
| | Version | 100 % | 55 % | 66 % | 90 % |
| | Total | 100 % | 66 % | 83 % | 94 % |

Data are mean ± SD.
n.s., no significantly above chance level (one-sample sign test, $p > 0.05$.).
